# Beyond Static Models: The Dynamic Interplay of Facial Emotions and Attentional Scope

**DOI:** 10.1101/2024.07.15.603432

**Authors:** Kesong Hu, Shuchang He, Qi Li, Chiang-Shan R. Li

## Abstract

The interplay between emotion and attention has long been intensely scrutinized, with competing theories proposing divergent mechanisms. Building on our previous work, here we present evidence that refines these perspectives, revealing a nuanced, temporally dynamic relationship between emotional stimuli and attentional focus. Using a modified Flanker task with facial emotion cues, we demonstrate that the effects of emotional stimuli on attention evolve over time, contrary to traditional fixed-effect assumptions. Our results show distinct temporal patterns: Neutral faces elicited typical flanker effects initially, but only interference persisted later. Early-stage happy faces amplified flanker facilitation but not interference, while threat faces augmented flanker interference but not facilitation. In the late stage, flanker facilitation disappeared across all emotion conditions, and interference patterns converged, mirroring the neutral face condition. These findings indicate emotion’s influence on attention is more complex and dynamic than previously recognized, potentially reflecting learning or habituation processes. We propose a new framework for understanding emotion-attention interactions that transcends traditional dichotomies of attention focus and approach-avoidance, offering a more nuanced perspective on this critical cognitive interface.

---

Attention, the ability to select and focus on relevant stimuli in our environment, is fundamental to perception, learning, and decision-making. However, this cognitive process is profoundly influenced by emotion (Anderson et al., 2011; Hu et al., 2018). While the interaction between emotion and attention is well-established, the precise mechanisms underlying this relationship remain a subject of debate. Two prominent frameworks have been proposed to explain the influence of emotion on attention. The attention focus theory posits that emotions tune perceptual systems to focus either broadly or narrowly (Fredrickson, 2004). According to this theory, positive affect leads to a broader attentional scope, while negative affect narrows it. Supporting evidence includes studies showing that manic individuals, experiencing elated moods, focus more broadly (Andreasen & Powers, 1975), and that positive affect can impair visual selective attention by increasing the processing of spatially adjacent flanking distractors (Rowe et al., 2007). But negative emotions, such as threat and stress, have been shown to narrow attentional focus (Easterbrook, 1959). Arousal during negative affective states is linked to a constriction of attentional focus, and habitually anxious and depressed individuals usually attend more to details (Basso et al., 1996). However, the evidence for the attention focus theory is not entirely conclusive (Gable & Harmon-jones, 2008; Harmon-jones et al., 2013; Pool et al., 2016; Vanlessen et al., 2016). An alternative perspective, the affect-as-information theory, rejects the fixed link between emotions and attentional scope(Clore & Huntsinger, 2009; Schwarz & Clore, 1983). This theory proposes that affective reactions are experienced in relation to whatever is in mind at the time (Clore et al., 2001; Schwarz & Clore, 1983). The magnitude of these effects varies as a function of arousal, occurring rapidly and showing significantly more prominent effects in initial measurements compared to later ones (Pool et al., 2016). The affect-as-information theory suggests that affective information simply provides the value of thoughts and responses at the moment (Clore & Huntsinger, 2009), resulting in a transient effect. This framework implies a more flexible influence of emotion on attentional scope. For instance, if a local focus is dominant, positive affect may narrow attentional scope, while negative affect may broaden it. Moreover, both positive and negative emotions could improve conflict processing(Kanske & Kotz, 2010, 2011; Li et al., 2014). This ongoing debate underscores the complex and dynamic nature of the relationship between emotion and attention, highlighting the need for further research to elucidate the underlying mechanisms and temporal dynamics of this interaction.

How facial expressions influence attention focus is a topic of great interest to scientists and laypeople alike. ***In this study***, we aimed to investigate how facial expressions of happiness and threat modulate the allocation of focused attention. Facial expressions convey a wide range of emotions, which are meaningful sources of social and biological information that can significantly affect attention, consciously and unconsciously. Converging evidence from behavioral and neuroimaging studies suggests that attention is preferentially “captured” by emotional facial expressions(Öhman et al., 2001; Schindler & Bublatzky, 2020). Several studies have used the Flanker task to assess the effects of facial emotions on focused attention (Barratt & Bundesen, 2012; Booy et al., 2023; Fenske & Eastwood, 2003; Horstmann et al., 2006; Schmidt & Schmidt, 2013; Tannert & Rothermund, 2020; Watson et al., 2012). In this paradigm, participants are presented with a row of stimuli (e.g., faces) with a target stimulus in the center and flanker stimuli on either side. The flankers can be either compatible (congruent trials) or incompatible (incongruent trials) with the target stimulus. Participants are instructed to respond to the target stimulus while ignoring the flankers (Eriksen & Eriksen, 1974; Gotlib & McCann, 1984). Previous research has shown that it is generally more difficult to suppress irrelevant information on the Flanker task when stimuli are affective. For instance, Fenske and Eastwood (2003) found that the flanker compatibility effect was smaller when target faces expressed negative compared to positive emotion or neutral targets. Tannert and Rothermund (2020) reported that the influences of face-induced expressions on attention depended on emotion being task-relevant or not, and attending to emotional faces was much less automatic than previously assumed. These studies have shed significant light on the intricate connection between facial emotion and attention. However, a common limitation in these studies is the use of faces as flanker stimuli. This approach inadvertently undermines the fundamental concept of investigating attention allocation to task-irrelevant flanker features when flanker effects are computed for different face targets. Moreover, whether and how facial emotion and attention interact across early and late stages remains unknown, leaving open the question of whether individuals exhibit habituation to the effects of facial emotion.

In addition, the current literature shows inconsistencies in the definition of flanker effects and data analysis (Ridderinkhof et al., 2021). Some studies have used the index “flanker compatibility,” calculated as the difference between response times (RTs) in trials with incongruent and congruent flankers (e.g., Fenske & Eastwood, 2003), while others directly termed this compatibility index as “flanker effects” (Mattler, 2003). This inconsistency is partly due to the absence of neutral flanker trials in these experiments. However, relying solely on this kind of index (i.e., RT_Incongruent_-RT_congruent_) to inform theories overlooks important behavioral information from neutral trials. Consequently, the individual effects of flanker facilitation (e.g., RT_neutral_-RT_congruent_) and interference (e.g., RT_Incongruent_-RT_neutral_) remain elusive. Our study aims to address these limitations and provide a more comprehensive understanding of how facial expressions modulate focused attention, taking into account both the nature of the stimuli and the temporal dynamics of the effect.

One approach to investigating the influence of facial emotion on attention focus involves eliciting an emotional state and subsequently exploring its impact on perceptual processing. For instance, Finucane employed film clips to induce fear and anger moods in participants, who then performed a feature-relevant flanker task(Finucane, 2011)^1^. This study reported that negative moods decreased flanker compatibility. Adopting a similar method, Wegbreit and colleagues evaluated how film-induced anxious and happy moods affected attention processing in the feature-relevant flanker task (Wegbreit et al., 2015). They found that negative, anxious moods generally narrowed the featural scope of visual attention, while positive moods had no discernible impact on attention in feature space. Due to the absence of neutral flanker trials, these authors employed flanker compatibility to present their data. **A central goal** of our study was to examine the extent to which spatial attention focus is modulated by preceding emotions, specifically happiness and threat. We investigated the effects of facial expressions inducing happiness and threat on selective attention and conflict processing, focusing on a more spatial-based process in the subsequent flanker task. We employed an arrow-flanker task, where participants were instructed to respond to the central arrow (the target, denoted as < or >) while disregarding extraneous flanker stimuli (e.g., << or >>). This task demands that participants selectively process the relevant element (the target arrow) while filtering out irrelevant input (the flanker arrows). In our study, participants were initially exposed to either a positive, negative, or neutral facial expression and subsequently completed the more spatial-based flanker task. We hypothesized that the mere presence of biologically salient stimuli, such as facial expressions, would influence subsequent feature processing. Using this paradigm, we aimed to address the limitations of previous studies and provide a more comprehensive understanding of how facial expressions modulate focused attention. This design enables us to explore whether individuals show habituation to the effects of facial emotion over time, a question that remains largely unexplored in the current literature.

Understanding the influences of emotion on flanker facilitation and interference can significantly contribute to evaluating affect-attention focus frameworks (Ekman et al., 2013). The *attention focus theory* posits that positive and negative emotions would broaden and narrow attention focus, respectively. As an extension, positive emotion was expected to result in heightened flanker facilitation as well as more pronounced flanker interference. Conversely, negative emotion was expected to yield reduced flanker facilitation and diminished flanker interference. By contrast, the *affect-as-information* theory hypothesized that emotions could either enhance or impair performance, contingent on the individual’s mindset or disposition. While traditional attention theories often portray attention allocation as a fixed and unwavering resource, one of our primary objectives in the present study was to investigate the dynamic nature of emotion’s influence on attention (Pourtois & Vuilleumier, 2006). This aligns with existing literature, which suggests that attending to emotional faces is far less automatic than previously assumed (Tannert & Rothermund, 2020), and that temporal change is the key to attentional capture (von Muhlenen et al., 2005). What sets our study apart is the utilization of a modified flanker task, featuring a cue positioned directly above the target stimulus, to assess the impact of emotion on attention focus (Li et al., 2023). Using that modified flanker task paradigm, we previously reported that both healthy and patient participants performed the task well, with healthy controls showing typical attention patterns where they were initially distracted by conflicting flanking arrows but learned to ignore them over time. This additional cue has the potential to either enhance performance by directing attention to its corresponding column or act as a distractor if emotions broaden participants’ attention. This approach allows us to explore the dynamic nature of emotional influences on attention, potentially revealing how these effects may change over the course of the experiment. We hypothesized that participants may adapt and utilize the cue over time to guide attentional orientation, leading to the effects of emotion becoming less or more pronounced in the later stages of the experiment.

## Methods

### Participants

Twenty-one participants (mean age = 28 years, SD = 1.60) took part in the experiment. This sample size aligns with our previous emotion and cognition studies (Hu et al., 2014; Hu et al., 2011, 2015; Hu, Rosa, et al., 2018; Li et al., 2023). Participants were undergraduate and graduate students from Peking University, Beijing. All were right-handed with normal or corrected-to-normal vision, and were naive to the study’s purpose. Written informed consent was obtained from all participants before the study began. The research followed the guidelines of the Helsinki Declaration and received approval from the local ethics committee.

### Apparatus and Procedure

We employed a modified Flanker task (Eriksen & Eriksen, 1974) as our experimental paradigm. The stimuli included arrays of five arrows, with a central target arrow (e.g., → or ←) flanked by two identical arrows on each side (e.g., →→, or ←←). Additionally, a cue (either a circle or a triangle) was presented directly above the central arrow stimulus.

**Figure 1.**
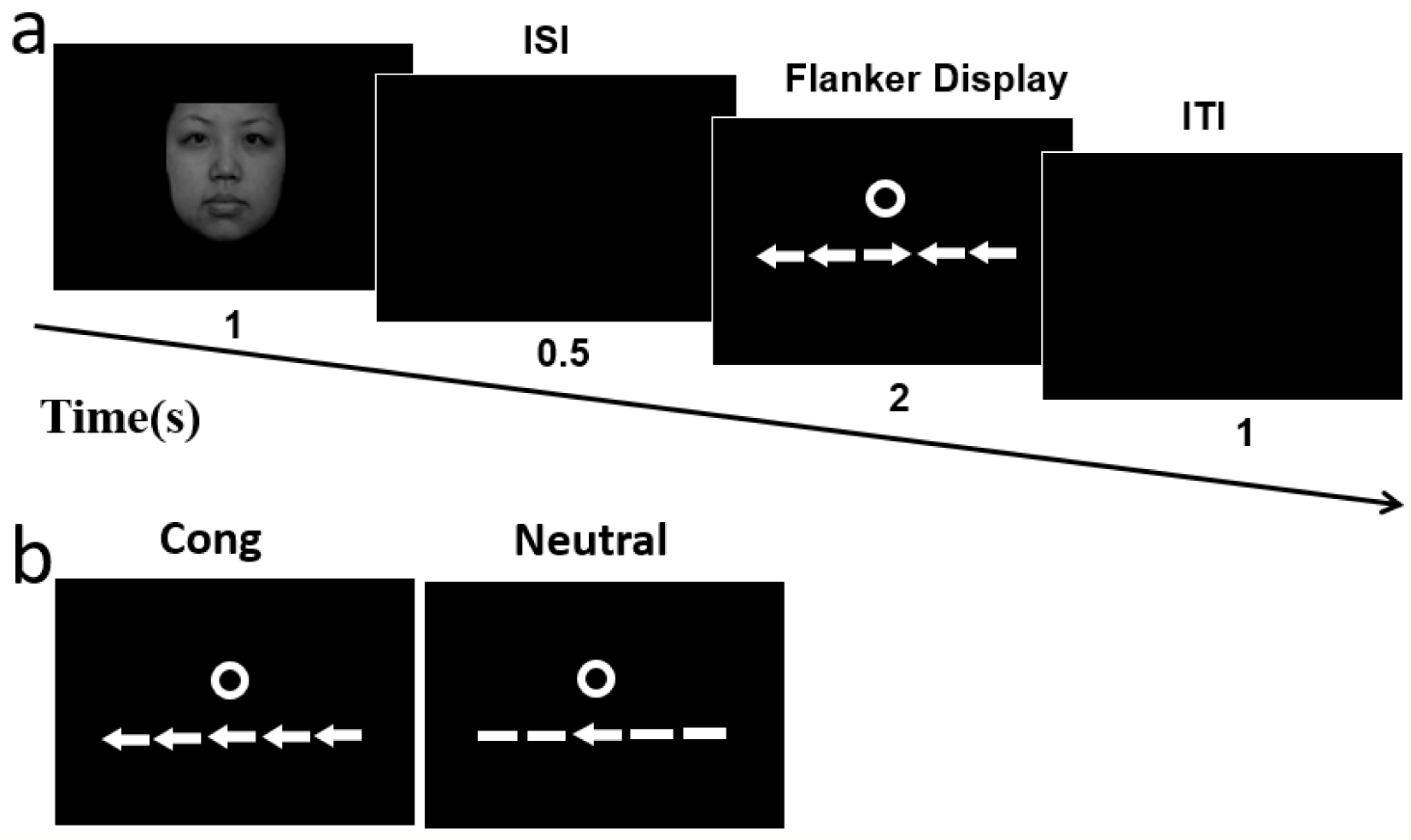
Task design and stimuli. (a) Trial sequence: A facial expression (1s) was followed by a fixation display (0.5s), then the target display (2s). The target display included a flanker stimulus (neutral, congruent, or incongruent) and a cue (circle or triangle) above the central arrow. Each trial ended with a 1sblank display. Note: Not all task phases are shown for simplicity. (b) Examples of congruent and neutral flanker displays. Facial expressions were categorized as happy, neutral, or threat. See main text for full details.

Figure 1 illustrates the sequence of events for a trial. Each trial began with a 1000-ms facial expression stimulus, followed by a 500-ms fixation display, and then a target display showing an array of five arrows on a black background for 2000 ms. The happy, neutral, and threatening facial stimuli were sourced from the Chinese Affective Picture System (CAPS) (Lu et al., 2005). Three types of trials were presented: congruent (e.g., →→→→→, or ←←←←←), incongruent (e.g., → →←→→, or ←←→←←), and neutral (e.g., ——→——, or ——←——). The arrays subtended 3.9° of visual angle (4.8 cm wide). An additional cue (1°, either a circle or a triangle) appeared above the central arrow simultaneously. Participants were instructed to respond as quickly and accurately as possible to indicate the direction of the central arrow while ignoring other stimuli. Each trial concluded with a 1000-ms blank screen. Responses were recorded using the NumLock keys on a computer keyboard placed in front of the participants. The inclusion of the additional cue above the central arrow in the flanker display made our study different from most previous Flanker task studies. This cue could potentially guide attention to the target, as it was aligned with the target stimulus. Participants might learn to use this cue to orient their attention over time, potentially reducing or eliminating flanker effects for both congruent and incongruent trials, particularly in later trials. However, the cue would only benefit performance if participants could effectively focus on this additional stimulus (Li et al., 2023). Upon completion, participants were debriefed about their experiences, feelings, and strategies used during the experiment.

Before the main experiment, participants completed a 30-trial practice session (unanalyzed). The experiment comprised six runs of 90 trials each, with short breaks between runs. Facial expression (happy, neutral, and threatening), cue type (circle or triangle), and congruency (congruent, neutral, and incongruent) factors were balanced. Trials were presented to ensure that no congruency condition was repeated in consecutive trials, minimizing potential priming effects (Davelaar & Stevens, 2009; Hu et al., 2012; Kesong Hu et al., 2015; Mayr et al., 2003; Verbruggen et al., 2006).

### Statistical Analyses

We analyzed correct response reaction times (RTs, in ms) and error rates separately. RTs exceeding ±3 SD from the condition-specific mean were excluded, resulting in the removal of 1.5% of all trials.

Our analysis was based on the following expectation: In the neutral emotion condition, we should observe standard flanker task effects, with faster RTs in congruent trials (facilitation effect) and slower RTs in incongruent trials (interference effect) compared to neutral trials. To test the difference between emotion conditions, we performed a repeated measure analysis of variance (ANOVA) on the RTs (based on correct responses) and error rates with the facial expression (happy, neutral, and threat) and congruency (congruent, neutral, and incongruent) as two within-subject factors. To explore the potential attentional adjustment across trials, we used our previous analysis approaches (Li et al., 2023). We ran ANOVAs to assess the flanker interference and facilitation effects on RTs across early and late trials for both control and emotional trials. Consistent with our previous work, we defined the early stage as Run 1 and the later stage as the mean data from Runs 2 to 6.

Additionally, we conducted planned independent t-tests to examine differences between conditions. Unless otherwise specified, the significance level for main statistical analyses was set at p < 0.05. When the sphericity assumption was violated in ANOVAs, we applied the Greenhouse– Geisser correction (Richard Jennings & Wood, 1976).

## Results

Research suggests that emotion and attention processing may be dynamic (Okon-Singer et al., 2013), and the influence of emotion on attention may be transient (Tannert & Rothermund, 2020; von Muhlenen et al., 2005). Consistent with our previous work (Li et al., 2023), we defined the early stage as Run 1 and the later stage as the mean data from Runs 2 to 6. We focused our analyses on reaction times (RTs) only, as the error rate was negligible (0.7%) and showed minimal variation across participants, precluding meaningful statistical analysis.

To validate our analysis approach, we first explored the stage effect using a repeated-measures ANOVA. This analysis included within-subject factors of Facial Emotion (happy, neutral, and threat), Congruency (congruent and neutral), and Stage (early vs. late) for RTs. The three-way interaction of Stage × Emotion × Congruency was significant, F(2, 40) = 5.18, p = .010, η² = .21, indicating a temporal effect on emotion and flanker facilitation, and supporting our stage-based analysis. We then conducted a further analysis using a 3 (Facial Emotion: happy, neutral, threat) × 2 (Congruency: neutral, incongruent) × 2 (Stage: early, late) repeated-measures ANOVA for RTs. This analysis also revealed a significant three-way interaction, F(2, 40) = 5.17, p = .010, η² = .21, further suggesting a temporal effect on emotion and flanker interference, and also validating our stage-based approach.

### Early stage: Flanker facilitation vs. interference

To assess the impact of emotion on attention focus in the early stage, we first examined whether typical flanker facilitation and interference effects were present in the neutral emotion condition (Fig. 2**, middle**). As expected, flanker facilitation was significant, t(20) = 2.44, p = .024, Cohen’s d = .531. Also, flanker interference was significant, t(20) = 4.65, p < .001, Cohen’s d = 1.015. We also compared the magnitudes of flanker interference and facilitation. Under neutral emotion conditions, flanker facilitation was significantly smaller than flanker interference (t(20) = 3.17, p = .005, Cohen’s d = .692), consistent with typical findings in the flanker task literature (Eriksen & Eriksen, 1974). Importantly, emotion did not significantly affect RTs in neutral flanker trials. We found no significant differences in RTs between happy and neutral emotions (t(20) = 1.08, p = .29, Cohen’s d = .236), or between threat and neutral emotions (t(20) = 1.59, p = .13, Cohen’s d = .346). Given these baseline findings, we conducted further analyses to explore the specific effects of emotion on congruent and incongruent trial processing.

**Figure 2.**
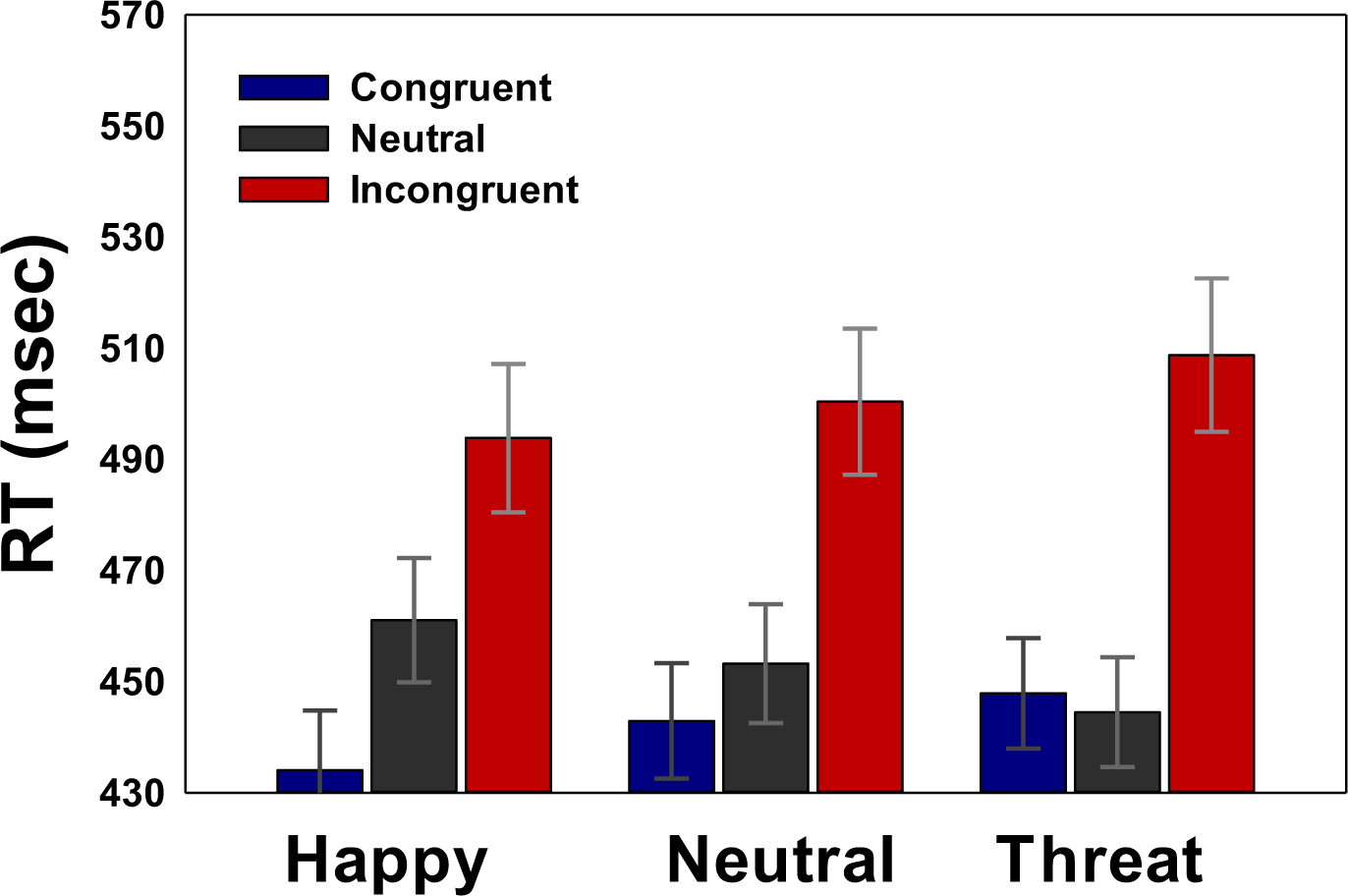
Reaction times (RTs) in the flanker task across congruent, neutral, and incongruent conditions, featuring happy, neutral, and threat facial expressions during the early stage.

### Flanker facilitation

We probed the facilitation effect with a 2 Congruency (neutral, congruent) x 3 Emotion (Happy, neutral, threat) repeated-measures ANOVA (Fig. 2). The main effect of Congruency was significant, F(1, 20) = 8.32, p = .009, η² = .294, suggesting slower RTs in neutral compared to congruent trials. The main effect of Emotion was non-significant, F(2, 40) = .06, p = .94, η² = .003. Critically, the Congruency × Emotion interaction was significant, F(2, 40) = 5.46, p = .008, η² = .214, suggesting that the RT difference between neutral and congruent trials varied across emotions.

The flanker facilitation effect is presented in Fig 3 (top panel). The magnitude of flanker facilitation was -3.40, 10.30, and 26.96 ms for threatening, neutral, and happy emotion conditions, respectively. The planned t-test showed that the flanker facilitation effect was significant in positive emotion conditions, t(20)=3.71, p=.001, Cohen’s d=.810. The facilitation effect was eliminated in threat emotion, t(20)=.44, p=.666, Cohen’s d=.096. In particular, the flanker facilitation was statistically larger in happy than neutral conditions. t(20)=2.09, p=.049, Cohen’d =.457. Furthermore, a significant linear trend was observed across happy, neutral, and threatening emotion conditions, F(1, 20) = 7.34, p = .014, η² = .268, suggesting a systematic change in facilitation effects across emotional valence.

**Figure 3.**
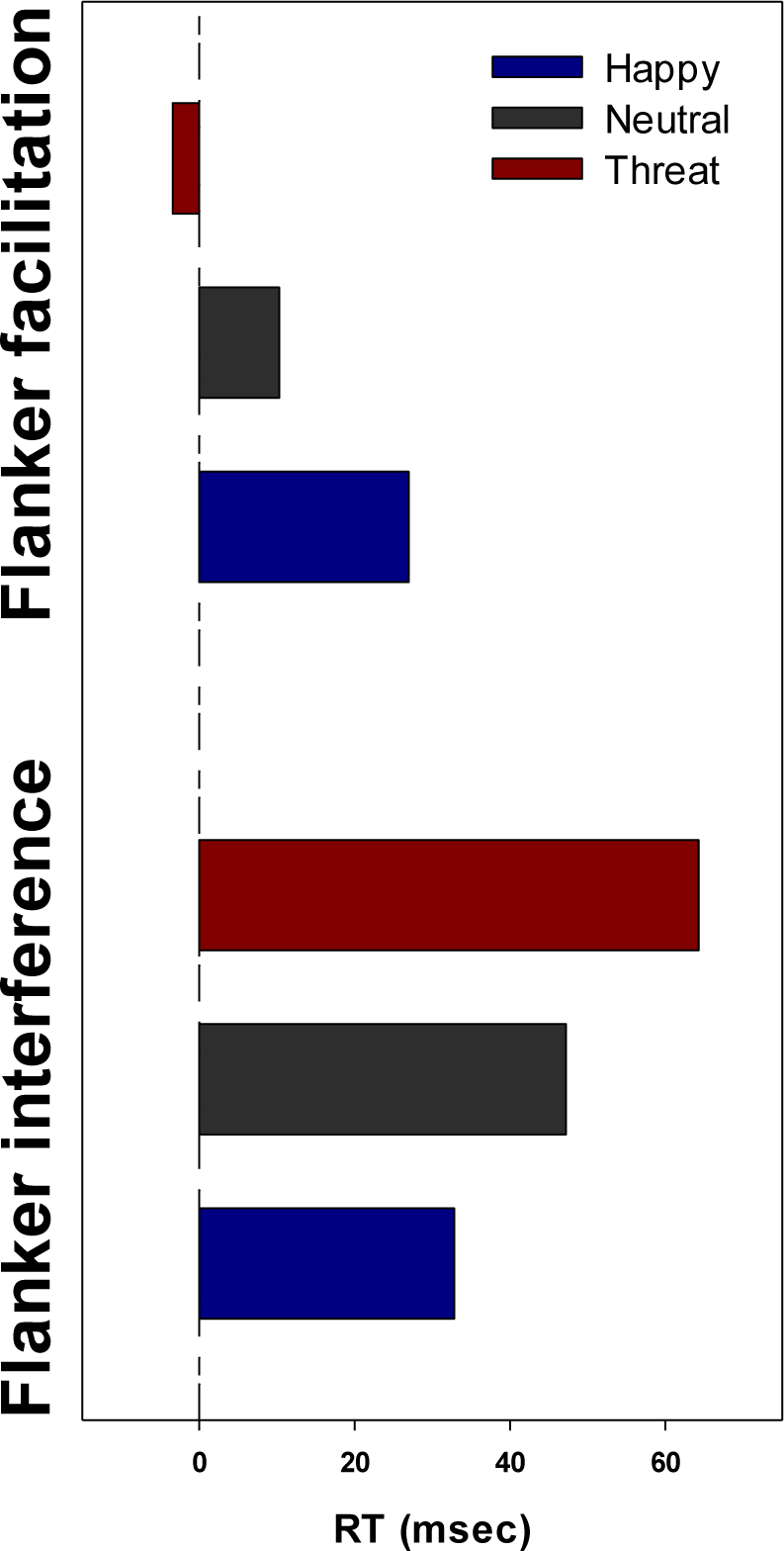
Flanker effects during the early stage. Top panel: Facilitation effects for happy, neutral, and threat facial expressions. Bottom panel: Interference effects for the same facial expression conditions.

### Flanker interference

Similarly, we examined the interference effect using a 2 Congruency (neutral, incongruent) x 3 Emotion (Happy, neutral, threat) repeated-measures ANOVA (Fig. 2). The main effect of Congruency was significant, F(1, 20) = 47.40, p < .0001, η² = .703, indicating slower RTs in incongruent compared to neutral trials. The main effect of Emotion was non-significant, F(2, 40) = .02, p = .98, η² = .001. Critically, the Congruency × Emotion interaction was significant, F(2, 40) = 3.78, p = .031, η² = .159, suggesting that the RT difference between incongruent and neutral trials varied across emotions.

The effect of flanker interference is presented in Fig 3 (bottom panel). The magnitude of flanker interference was 64.27, 47.18, and 32.82 ms for threatening, neutral, and happy emotion conditions, respectively. The planned t-test showed that the flanker interference effects were all significant across the emotion conditions, with the smallest effect being t(20) = 3.31, p = .004, Cohen’s d = .722. The trend effect across positive, neutral, and threat emotions was significant, F(1,20)=11.51, p=.003, η² = .365. The flanker interference was larger in threat than happy emotion condition (size =31.5 ms), t(20)=3.39, p=.003, Cohen’s d =.740. Collectively, these results demonstrate that happy facial expressions enhanced flanker facilitation but not interference, while threatening facial expressions enhanced flanker interference but not facilitation. This pattern suggests a dissociation in how positive and negative emotions modulate different aspects of attentional control.

### Later stage: Flanker facilitation vs. interference

#### Flanker facilitation

In the later stage, we observed no facilitation effects across the three emotion conditions (Fig. 4). This was confirmed with a 2 Congruency (neutral, congruent) x 3 Emotion (Happy, neutral, threat) repeated-measures ANOVA. The Congruency × Emotion interaction was not significant (F(2, 40) = .189, p = .83, η² = .009). Additionally, neither the main effect of Congruency (F(1, 20) = .84, p = .37, η² = .040) nor the main effect of Emotion (F(2, 31) = .19, p = .83, η² = .009) reached significance. The absence of facilitation effects in the later stage suggests an attentional adjustment occurred during the course of the experiment.

**Figure 4.**
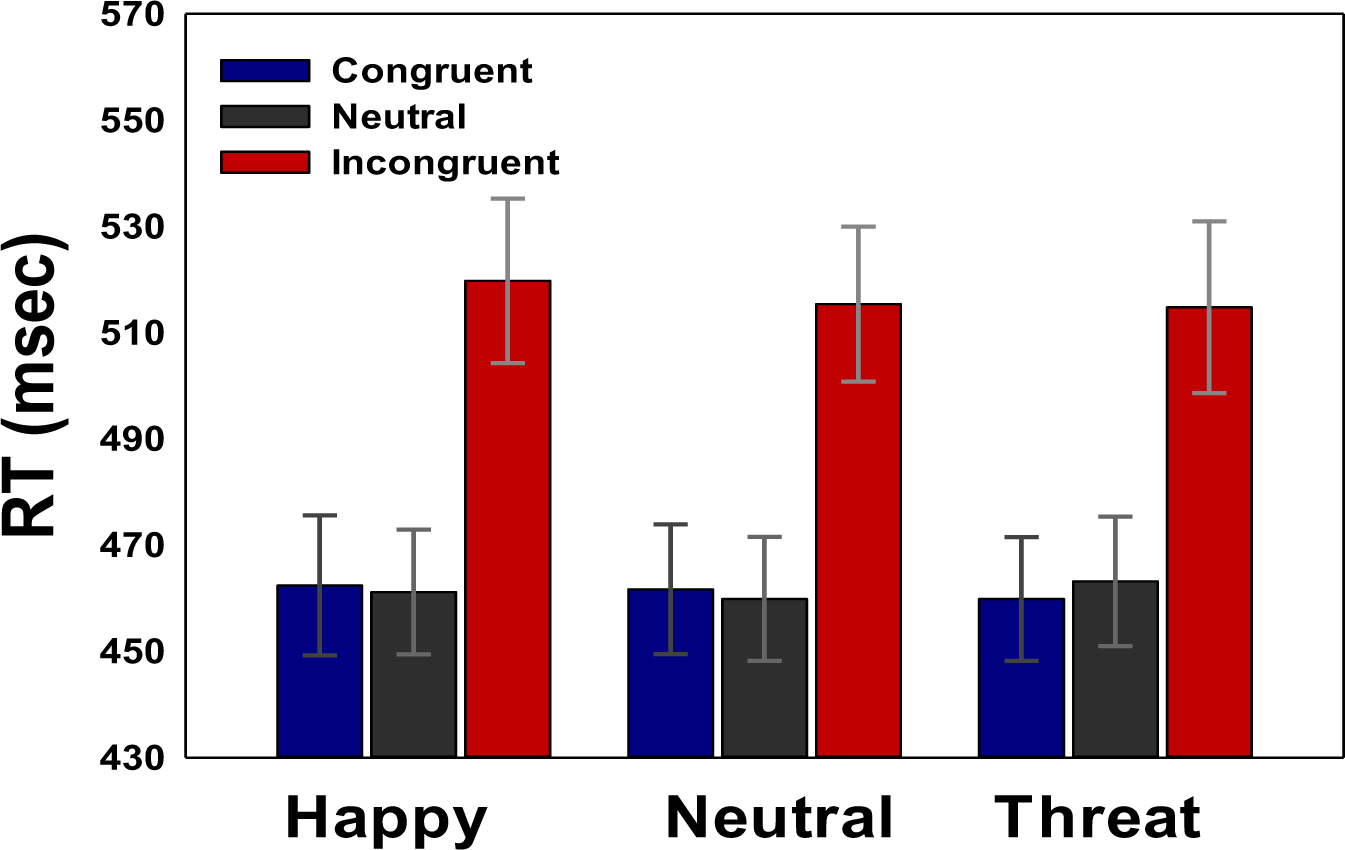
Reaction times (RTs) in the flanker task during the late stage, comparing congruent, neutral, and incongruent conditions across happy, neutral, and threat facial expressions.

#### Flanker interference

We examined the interference effect in the later stage using a 2 Congruency (neutral, incongruent) x 3 Emotion (Happy, neutral, threat) repeated-measures ANOVA (Fig. 4). The main effect of Congruency was significant, F(1, 20) = 50.68, p < .0001, η² = .717, indicating slower RTs in incongruent compared to neutral trials. Emotion had no significant main effect, F(2, 40) = .96, p = .39, η² = .046. The Congruency × Emotion interaction was also non-significant, F(2, 40) = 1.29, p = .286, η² = .061, suggesting that the RT difference between incongruent and neutral trials was consistent across emotions. These results indicate a shift in attentional processing from the early to the later stage, with the disappearance of facilitation effects and the persistence of interference effects across all emotion conditions.

### Comparison: Early vs. Late Stage

To summarize, we compared the flanker interference effects between the late and early stages^2^(Fig.5**, bottom panel**). The change in flanker interference between the late and early stages in the neutral emotion condition was not significant, t(20) = 0.789, p = .439, Cohen’s d = .172. However, flanker interference was significantly larger in the late than in the early stage for the happy condition, t(20) = 2.69, p = .007, Cohen’s d = .587. This pattern was reversed in the threatening emotion condition, t(20) = 1.94, p = .034, Cohen’s d = .423.

**Figure 5.**
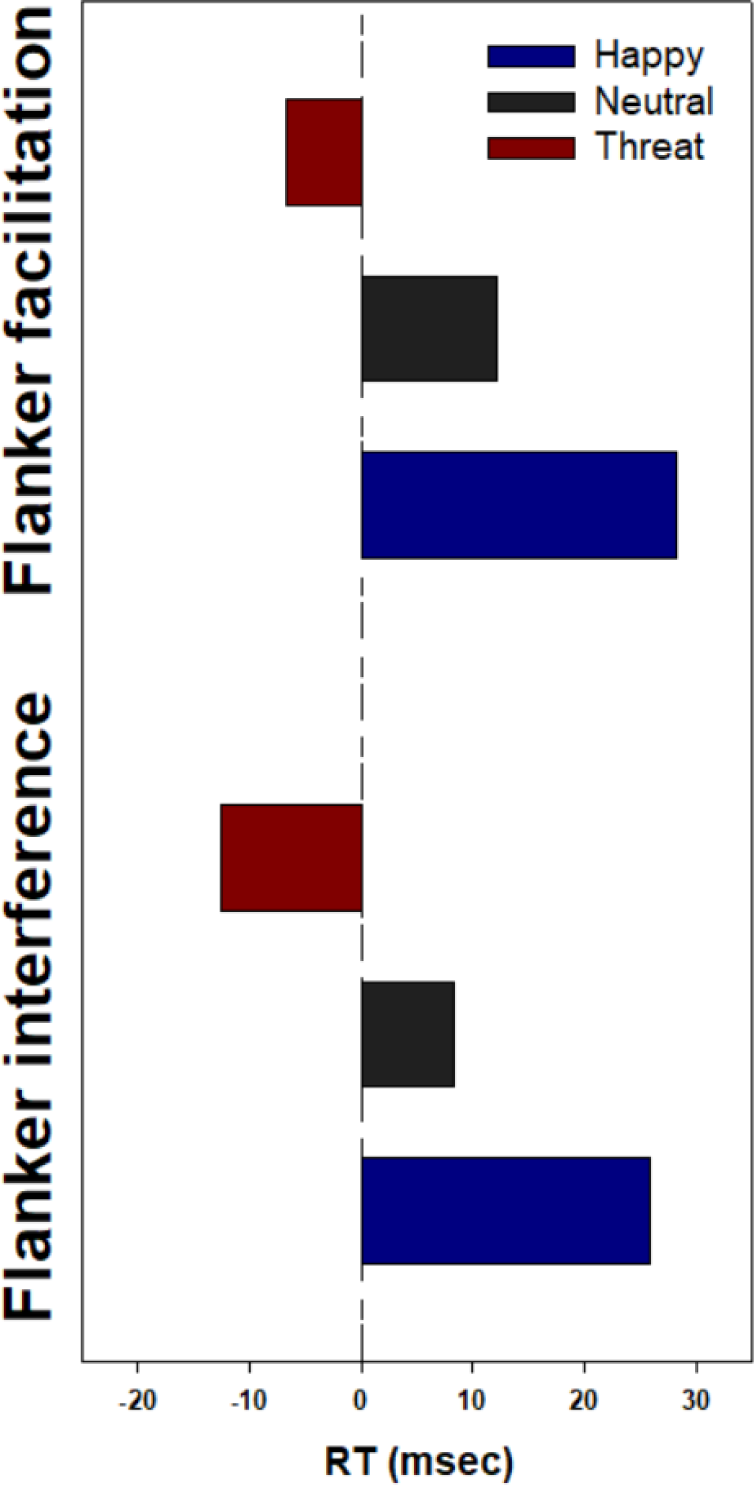
Comparison of flanker effects across emotional conditions (happy, neutral, and threat) in early and late stages. Top panel: Facilitation effects contrasting early and late stages. Bottom panel: Interference effects contrasting late and early stages.

Although flanker facilitation was only present in the early stage, we conducted a parallel comparison for completeness (Fig.5**, top panel**). The index of flanker facilitation was more prominent in the early than the late stage for the neutral emotion condition, t(20) = 1.96, p = .033, Cohen’s d = .427. This effect was more pronounced in the happy emotion condition, t(20) = 3.09, p = .003, Cohen’s d = .675. Under the threatening condition, flanker facilitation was absent in both early and late stages, with no significant difference between stages, t(20) = .911, p = .187, Cohen’s d = .199. These comparisons highlight the dynamic nature of emotional influences on attention, with different patterns of change for facilitation and interference effects across emotional conditions and stages of the task.

## Discussion

Building on our previous research (Li et al., 2023), this study investigates the dynamic interplay between emotion and attention using a modified Flanker task. We examined how facial expressions of happiness and threat modulate focused attention over time. Our findings reveal that facial expressions influence attention focus differently depending on congruency and task stage. In early trials, happy faces enhanced flanker facilitation, while threatening faces amplified interference. Crucially, these emotion-specific effects dissipated in later trials. Flanker facilitation disappeared across all emotion conditions, while interference persisted but didn’t differ between happy and threat conditions. These results show the influence of facial expressions on attention focus evolves throughout the task, challenging assumptions about sustained emotional effects on attention. This temporal shift suggests a complex, dynamic relationship between emotion and attention, which we discuss in detail below.

Firstly, in the initial phase of the neutral emotion condition, we saw both flanker facilitation and interference, consistent with established findings in the literature (Eriksen & Eriksen, 1974). Reaction times (RTs) were shorter for congruent flankers (e.g., →→→→→, or ←←←←←) compared to neutral flankers (e.g., ——→——, or ——←——), suggesting significant flanker facilitation. Conversely, RTs were longer for incongruent flankers (e.g., →→←→→, or ←←→←←) compared to neutral flankers, suggesting significant flanker interference. Flanker compatibility was also evident, with longer RTs for incongruent flankers compared to congruent ones. Notably, the flanker facilitation effect was smaller than the flanker interference effect. Interestingly, in the later stage of the experiment, flanker facilitation vanished while flanker interference persisted. This pattern suggests a dynamic adjustment of participants’ attention over the course of the trials. As the experiment progressed, participants might have realized that the additional cue could serve as a clue to the target or target column, or they might have learned to ignore it effectively. Both strategies resulted in a narrowing of attention focus in the later stage, as confirmed during debriefing. It’s worth noting that flanker facilitation is more susceptible to changes in the perceptual setting, making it a more delicate effect. This sensitivity likely contributed to the observed temporal effect on congruent trials. In contrast, flanker interference appears to be more robust and less likely to be modulated by the cue. Some researchers have suggested that flanker interference might occur independently of spatial attention (Ro et al., 2002), and the facilitation effect is neither dependent on the nature of interference nor the amount of cognitive control engaged in the task(Davranche et al., 2009). Collectively, our findings demonstrate that the commonly observed flanker effects are temporal in nature, and that participants exhibited attentional adjustment with this modified arrow-flanker task. This underscores that an individual’s attention focus is not constant but fluctuates over time. The focus of attention is contingent upon the perceptual set, and the magnitude of this influence varies across distinct time windows. These results highlight the dynamic nature of attention allocation and the importance of considering temporal factors in understanding attentional processes.

Our primary aim was to investigate how facial emotions influence attention focus, a process that is fundamentally bottom-up. Attention can be broadly categorized into two types: rule selection and feature input selection. While the Stroop task is widely used to investigate the influence of emotion on attention (Hu et al., 2012; Kesong Hu et al., 2015; Stroop, 1935), its performance relies on the application of specific rules, such as distinguishing between ink color and word color. The competition between these rules requires more than simple selection, introducing additional complexity to the task. In contrast, our Flanker task required participants to respond to the central arrow (the target, denoted as < or >) while disregarding extraneous flanker stimuli (e.g., << or >>). This design allowed us to assess the impact of emotions on spatial input selection without the complexities introduced by rule selection. Specifically, this arrow Flanker task is considered a spatial-based task due to the inherent spatial directional meaning conveyed by the arrows (Ridderinkhof et al., 2021). The spatial nature of the task also aligns well with the spatial aspects of attention, potentially offering insights into how emotions might differentially affect various spatial dimensions of attentional focus.

Secondly, our study revealed that positive and negative emotions differentially modulated attentional focus. The main part of this study was to assess the interaction of emotion and attention, and to evaluate current theories given our findings. In the early stage, RTs for neutral flanker trials (e.g., ——→——, or ——←——) did not differ across happy, neutral, and threat facial expression conditions. However, compared to the typical flanker facilitation observed in the neutral condition, happy facial expressions amplified flanker facilitation, while threat facial expressions eliminated it. This suggests that happy facial expressions broaden the scope of attention, while threat facial expressions narrow it, as reflected in the flanker facilitation effect. These findings align with the *attention focus theory*, which posits that arousal and valence determine the breadth of attention. Contrary to the facilitation effects, threat facial emotions amplified flanker interference, while happy facial emotions decreased it. In other words, happiness improved performance, while threat impaired or inhibited it. These findings contradict the *attention focus theory*, which predicts that happy emotions should broaden the scope of attention (leading to increased flanker interference), and threat emotions should narrow it (resulting in decreased flanker interference). However, these results partially support the *affect-as-information theory,* which suggests that emotions can either enhance or impair performance depending on the individual’s mindset or disposition. Our findings both align with and extend previous research in this area. Fenske and Eastwood (2003) reported that the flanker compatibility effect was smaller in negative than in positive emotion or neutral conditions when using schematic facial expressions. Our work furthers this by demonstrating that negative emotion led to more attention allocation in incongruent processing, while positive emotion led to more attention allocation in congruent processing. These results are partially consistent with current literature. For instance, both positive and negative emotions have been reported to facilitate conflict processing (Kanske & Kotz, 2010, 2011). More specifically, our findings align with a behavioral and ERP study, which showed that negative emotion led to increased attention allocation on incongruent trials, while positive emotion led to increased attention allocation on congruent trials (Li et al., 2014). These results provide a nuanced understanding of how emotions modulate attention, suggesting that the effects are more complex than previously theorized and depend on the specific type of attentional process being engaged.

In the later stage of our experiment, we observed a striking shift in attentional patterns. Flanker facilitation disappeared across all emotion conditions, while flanker interference showed no significant differences between happy and threat conditions, mirroring the effect seen with neutral faces. These findings challenge the simplistic view that emotions consistently tune perceptual systems to focus either broadly or narrowly. While our data partially align with the *affect-as-information theory*, which proposes that emotions can either enhance or impair performance depending on an individual’s mindset or disposition, this theory fails to account for the crucial temporal factor we observed. Our results underscore the importance of considering time in emotion-attention interactions, a concept rooted in Posner’s seminal work (Posner, 1978). Posner proposed that visual information accumulates over time, with observers accessing this information at different points. Our findings are consistent with a meta-analysis by Pool and colleagues (Pool et al., 2016), which revealed that emotional effects on attention are more pronounced in early stages rather than later stages of processing. Our temporal analysis extends this understanding, demonstrating that the influence of emotion on attentional scope is dynamic rather than static. In the early stage of our experiment, positive and negative emotions differentially influenced performance on congruent and incongruent flanker trials, aligning to some extent with current theories. However, our data reveal this emotional influence on attention focus diminishes over time, likely as individuals habituate to the emotional stimuli and environment. Comparing early and late stages, we found that: 1) The flanker facilitation difference was more prominent in the happy than neutral emotion condition for congruent trials, but this effect was absent in the threat condition. 2) The flanker interference difference (late vs. early) was larger in the happy compared to neutral emotion condition for incongruent trials. Interestingly, this pattern was reversed in the threat condition compared to the neutral condition. These data suggest that the emotional influence on spatial attention focus is contingent on both the temporal factor (early and late) and task perceptual set (Huntsinger, 2012; Tannert & Rothermund, 2020). Our findings underscore the complex, dynamic nature of emotion-attention interactions and highlight the importance of considering temporal dynamics in understanding these processes.

Our findings seem to reveal that feelings and motivations fluctuate over time, and their impact on the temporal window of spatial attention focus also varies. This dynamic nature of emotional attention requires a refinement of current theories. The inextricable link between space and time, evident in both the physical world and the psychological realm (Hawking & Penrose, 2010; Jones, 1976; Large & Jones, 1999), underscores the fact that all life evolves across these dimensions (Kuppens & Verduyn, 2017). We propose a perspective that moves beyond a strict attention focus and approach-avoidance dichotomy, challenging longstanding claims about the sustained effects of emotion on attention. At a fundamental level, emotional and motivational influences exhibit two key aspects: 1) An inertia part: an inherent resistance to change; 2) A regulatory part: continuous adjustment to maximize alignment with the current desired state. Ultimately, these two aspects interact and determine how individuals’ emotions unfold in their influence on attention focus over time. Behaviorally, emotional stimuli can capture attention early in processing, but this attentional capture can evolve, fluctuate, and diminish over time. In the brain, this is likely due to its ability to adjust and habituate to emotional stimuli, resulting in changes in its influence on attention focus. The effect of emotion on attention focus is not static, and its effect goes down over time (Pourtois & Vuilleumier, 2006; von Muhlenen et al., 2005). Until the temporal factor is considered, our understanding of how facial expressions influence or determine our attention focus will remain incomplete (von Muhlenen et al., 2005). In essence, the influence of emotion on attention focus is a complex and dynamic phenomenon. Future research and theories must not only account for temporal dynamics but also consider the roles of perceptual sets and task requirements. By adopting this more nuanced approach, we can develop a more comprehensive understanding of how emotions shape our attentional processes over time, leading to more accurate predictions and interventions in both clinical and non-clinical settings.

Our evaluation of the temporal effects in the interaction between emotion and cognition was facilitated, in part, by our unique experimental design. We utilized a modified Flanker task that incorporated an additional cue positioned directly above the center of the display. This setup differs significantly from previous studies, allowing us to capture temporal dynamics that might otherwise go unnoticed. For instance, Wegbreit and colleagues (2015) employed a feature-based Flanker task (i.e., letter version) where flankers were occasionally colored differently than the target stimuli. In their setup, the “additional cue” was a feature of the target itself, potentially limiting participants’ ability to capitalize on it for task processing. Their findings indicated that an anxious mood led participants to focus on irrelevant features, while a positive mood did not influence attention in feature space(Wegbreit et al., 2015). Our study featured a task-irrelevant cue that could serve as a guide for directing attention to the central arrow stimulus. To appreciate this cue (otherwise considered noise), participants needed time. Some may have gradually realized the cue could be inhibited, triggering attentional regularities in the environment (Rhodes & Di Luca, 2016). This process aligns with Bayesian models of perception and cognition. Participants likely started with a prior in the early stage and then employed additional evidence to update this prior, refining their performance in the late stage(Barlow, 2001; Knill & Pouget, 2004). These dynamics culminated in an attention-emotion adjustment, evident in the neutral emotion condition where flanker facilitation was absent in the late stage. It’s worth noting that the facilitation effect is not contingent on the nature of interference or the level of cognitive control engaged in the task (Davranche et al., 2009). Relative to interference, the facilitation effect is generally smaller and more delicate in executive tasks, including both Flanker and Stroop tasks (Eriksen & Eriksen, 1974; Kesong Hu et al., 2015; Kalanthroff & Henik, 2013; MacLeod & MacDonald, 2000). This characteristic may have contributed to our ability to reveal the temporal effect on flanker effects.

Note that, unlike previous studies (Barratt & Bundesen, 2012; Booy et al., 2023; Fenske & Eastwood, 2003; Horstmann et al., 2006; Kuppens & Verduyn, 2015; Schmidt & Schmidt, 2013; Tannert & Rothermund, 2020; Watson et al., 2012), our work reveals the intricate interplay between positive and negative facial emotions and attention allocation dynamics, encompassing two key aspects. ***First***, we chose an arrow-based Flanker task over the traditional letter-based version. Arrows not only carry symbolic information but also provide spatial cues. As arrow directions are universally understood and learned early in life, they automatically convey spatial meaning (Mattler, 2003; Ridderinkhof et al., 2021). We categorize the letter Flanker task as feature-based, and the arrow Flanker task as spatial-based due to the inherent directional meaning conveyed. Our findings shed light on how facial expressions impact spatial attention, delving into a sequential process and building upon prior research on feature attention processing. ***Second***, unlike earlier studies (e.g., Fenske & Eastwood, 2003; Tannert & Rothermund, 2020), our study uniquely separated facial emotion and flanker stimuli during trials. This extension of prior research digs into the fundamental process, considering humans’ specialization in perceiving facial expressions. We used real rather than schematic faces as the latter lacks ecological value (Horstmann & Bauland, 2006). The manipulation of facial expressions within our experiment provides an ecologically valid approach for assessing emotion, attention, and model selection. By separating emotional stimuli from the attention task itself and using ecologically valid facial expressions, we can more accurately capture the dynamic nature of emotion-attention interactions as they unfold in real-world scenarios.

In conclusion, our findings demonstrate that facial expressions do not universally influence the scope of attention (Huntsinger, 2012, 2013). Instead, the interplay between emotion and attention focus is dynamic and exhibits temporal variations. This discovery lays the foundation for a new theoretical framework that incorporates the crucial yet often overlooked temporal factor in understanding the complexity of emotion-attention interactions. Emotions play a pivotal role in recruiting control mechanisms (Inzlicht et al., 2015), temporarily influencing whether individuals act on transient tendencies to focus or narrow their attentional windows(Huntsinger, 2012, 2013; Okon-Singer et al., 2013). Our research proposes a perspective that transcends the rigid attention focus and approach-avoidance dichotomy traditionally used in this field. Furthermore, we suggest that arousal and valence, in conjunction with the task itself, provide inputs to control-related processes that ultimately determine performance (Anderson, 2005; Dignath et al., 2020). These factors hold significant promise for future advancements in the fields of emotion and motivation science. Moving forward, several key areas justify further investigation: 1) Individual differences: Future research should explore how personal characteristics modulate the dynamic interaction between emotion and attention; 2) Spatial independence: Studies should aim to determine whether these effects manifest independently of spatial attention (Ro et al., 2002; Rowe et al., 2007); 3) Clinical implications: Investigating the potentially maladaptive influence of emotion on attention focus in patients with clinical conditions (Heylen et al., 2016; Hu et al., 2014) could provide valuable insights for both theoretical understanding and practical interventions. This perspective emphasizes the need for dynamic models that account for the temporal evolution of emotion-attention interactions, potentially leading to more effective interventions in clinical settings and a deeper understanding of human cognition in everyday life.

## Data Availability

The data are available from the corresponding authors upon request.

## Acknowledgements

We would like to express our gratitude to all participants for their contributions. We also acknowledge that all participants have provided their consent to publish the findings of this study.

## Footnotes

1 The letter flanker task presents a central target letter (e.g., H) flanked on either side by response-compatible (e.g., HHHHH) or response-incompatible (e.g., NNHNN) letters. This task is typically considered feature-based, in contrast to the arrow flanker task, which is generally regarded as spatial-based due to the inherent directional meaning conveyed by arrows. For further discussion, see (Ridderinkhof et al., 2021).

2 To explore trend changes, we used one-tailed planned t-tests to assess the emotion effect on attention.

